# Efficient long single molecule sequencing for cost effective and accurate sequencing, haplotyping, and *de novo* assembly

**DOI:** 10.1101/324392

**Authors:** Ou Wang, Robert Chin, Xiaofang Cheng, Michelle Ka Wu, Qing Mao, Jingbo Tang, Yuhui Sun, Ellis Anderson, Han K. Lam, Dan Chen, Yujun Zhou, Linying Wang, Fei Fan, Yan Zou, Yinlong Xie, Rebecca Yu Zhang, Snezana Drmanac, Darlene Nguyen, Chongjun Xu, Christian Villarosa, Scott Gablenz, Nina Barua, Staci Nguyen, Wenlan Tian, Jia Sophie Liu, Jingwan Wang, Xiao Liu, Xiaojuan Qi, Ao Chen, He Wang, Yuliang Dong, Wenwei Zhang, Andrei Alexeev, Huanming Yang, Jian Wang, Karsten Kristiansen, Xun Xu, Radoje Drmanac, Brock A. Peters

## Abstract

Obtaining accurate sequences from long DNA molecules is very important for genome assembly and other applications. Here we describe single tube long fragment read (stLFR), a technology that enables this a low cost. It is based on adding the same barcode sequence to sub-fragments of the original long DNA molecule (DNA co-barcoding). To achieve this efficiently, stLFR uses the surface of microbeads to create millions of miniaturized barcoding reactions in a single tube. Using a combinatorial process up to 3.6 billion unique barcode sequences were generated on beads, enabling practically non-redundant co-barcoding with 50 million barcodes per sample. Using stLFR, we demonstrate efficient unique co-barcoding of over 8 million 20-300 kb genomic DNA fragments. Analysis of the genome of the human genome NA12878 with stLFR demonstrated high quality variant calling and phasing into contigs up to N50 34 Mb. We also demonstrate detection of complex structural variants and complete diploid *de novo* assembly of NA12878. These analyses were all performed using single stLFR libraries and their construction did not significantly add to the time or cost of whole genome sequencing (WGS) library preparation. stLFR represents an easily automatable solution that enables high quality sequencing, phasing, SV detection, scaffolding, cost-effective diploid *de novo* genome assembly, and other long DNA sequencing applications.

---

To date, the vast majority of individual higher organism whole genome sequences lack information regarding the order of single to multi-base variants transmitted as contiguous blocks on homologous chromosomes, typically referred to as haplotypes. In addition, most sequenced genomes leave unresolved novel sequence not found in reference genomes, large structural variations, repeat sequences, and other regions which are difficult to analyze with current technologies. For many early genome studies this information was not critical and was overlooked. However, as we continue to move towards sequencing all species and a more complete understanding of how an individual’s genome contributes to the myriad phenotypes they exhibit, this missing information will become necessary. In order to achieve this a near “perfect genome” (Peters et al. 2014), that is an individual genome in which all bases are properly sequenced, homologous chromosomes are independently assembled, and almost no errors are made, will be required. Furthermore, this has to be achieved at low cost to be affordable for the billions of people around the world that will eventually be sequenced.

Numerous technologies, including direct single molecule sequencing (Levene et al. 2003; Zhang et al. 2006; Ma et al. 2010; Olasagasti et al. 2010; Fan et al. 2011; Kitzman et al. 2011; Suk et al. 2011; Duitama et al. 2012; Peters et al. 2012; Selvaraj et al. 2013; Amini et al. 2014; Kuleshov et al. 2014; Zheng et al. 2016), have recently been developed to generate at least some of this information. Most are based on the process of co-barcoding (Peters et al. 2014), that is, the addition of the same barcode to the sub-fragments of single long genomic DNA molecules. After sequencing the barcode information can be used to determine which reads are derived from the original long DNA molecule. This process was first described by Drmanac (Drmanac 2006) and implemented as a 384-well plate assay by Peters et al. (Peters et al. 2012). These approaches have been technically challenging to implement, are expensive, have lower data quality, do not analyze individual DNA molecules separately (*i.e.*, do not provide unique co-barcoding), or some combination of all four. In practice, most require a separate whole genome sequence to be generated by standard methods to improve variant calling. In addition, most can only provide haplotype information, but are unable to provide the other additional information necessary for perfect genome sequencing.

## Results

### stLFR library process

Here we describe implementation of stLFR technology (Drmanac 2013), an efficient approach for DNA co-barcoding with millions of barcodes enabled in a single tube. This is achieved by using the surface of a microbead as a replacement for a compartment (e.g., the well of a 384-well plate). Each bead carries many copies of a unique barcode sequence which is transferred to the sub-fragments of each long DNA molecule. These co-barcoded sub-fragments are then analyzed on common short read sequencing devices such as the BGISEQ-500 or equivalent. In our implementation of this approach we use a ligation-based combinatorial barcode generation strategy to create over 3.6 billion different barcodes in three ligation steps. For a single sample we use ~10-50 million of these barcoded beads to capture ~10-100 million long DNA molecules in a single tube. It is infrequent that two beads will share the same barcode because we sample 10-50 million beads from such a large library of total barcodes. Furthermore, in the case of using 50 million beads and 10 million long genomic DNA fragments, the vast majority of sub-fragments from each long DNA fragment are co-barcoded by a unique barcode. This is analogous to long-read single molecule sequencing and potentially enables powerful informatics approaches for *de novo* assembly. A similar but informatically limited and less efficient approach using only ~150,000 barcodes was recently described by Zhang et al. (Zhang et al. 2017). Importantly, stLFR is simple to perform and can be implemented with a relatively small investment in oligonucleotides to generate barcoded beads. Further, stLFR uses standard equipment found in most molecular biology laboratories and is sequencing technology agnostic. Finally, stLFR replaces standard NGS library preparation methods, requires only 1 ng of DNA, and does not add significantly to the cost of whole genome or whole exome library preparation with a total cost per sample of less than 30 dollars (Table 1).

**Table 1.** stLFR equipment and reagent cost (USD)

| Equipment | One-time cost | Per sample |
| --- | --- | --- |
| Sample rotator | ~500 |  |
| Incubator | ~2,000 |  |
| Magnetic separation rack | ~600 |  |
| Thermocycler | ~10,000 |  |
| **Reagents** |  |  |
| Barcode oligos | ~50,000<sup>1</sup> | 0.13 |
| Streptavidin labeled beads |  | 7 |
| Enzymes for barcoded bead construction |  | 4.40 |
| Enzymes for stLFR library construction |  | 17 |
| Total | ~63,100 | 28.53 |

<sup>1</sup>Barcode oligonucleotides are listed as a one-time cost because they cannot be purchased on a per sample basis. At a 100 nmol scale synthesis, the cost per sample of oligos is approximately 0.14 dollars.

The first step in stLFR is the insertion of a hybridization sequence at regular intervals along genomic DNA fragments. This is achieved through the incorporation of DNA sequences, by the Tn5 transposase, containing a single stranded region for hybridization and a double stranded sequence that is recognized by the enzyme and enables the transposition reaction (Figure 1a). Importantly, this step is done in solution, as opposed to having the insertion sequence linked directly to the bead (Zhang et al. 2017). This enables a very efficient incorporation of the hybridization sequence along the genomic DNA molecules. As previously observed (Amini et al. 2014), the transposase enzyme has the property of remaining bound to genomic DNA after the transposition event, effectively leaving the transposon-integrated long genomic DNA molecule intact. After the DNA has been treated with Tn5 it is diluted in hybridization buffer and added to 50 million ~2.8 um clonally barcoded beads in hybridization buffer. Each bead contains approximately 400,000 capture adapters, each containing the same barcode sequence. A portion of the capture adapter contains uracil nucleotides to enable destruction of unused adaptors in a later step. The mix is incubated under optimized temperature and buffer conditions during which time the transposon inserted DNA is captured to beads via the hybridization sequence. It has been suggested that genomic DNA in solution forms balls with both tails sticking out (Jo et al. 2009). This may enable the capture of long DNA fragments towards one end of the molecule followed by a rolling motion that wraps the genomic DNA molecule around the bead. Approximately every 7.8 nm on the surface of each bead there is a capture oligo. This enables a very uniform and high rate of sub-fragment capture. A 100 kb genomic fragment would wrap around a 2.8 um bead approximately 3 times. In our data, 300 kb is the longest fragment size captured suggesting larger beads may be necessary to capture longer DNA molecules. Beads are next collected and individual barcode sequences are transferred to each sub-fragment through ligation of the nick between the hybridization sequence and the capture adapter (Figure 1a). At this point the DNA/transposase complexes are disrupted producing sub-fragments less than 1 kb in size. Due to the large number of beads and high density of capture oligos per bead, the amount of excess adapter is four orders of magnitude greater than the amount of product. This huge unused adapter can overwhelm the following steps. In order to avoid this, we designed beads with capture oligos connected by the 5’ terminus. This enabled an exonuclease strategy to be developed that specifically degraded excess unused capture adapter.

**Figure 1.**
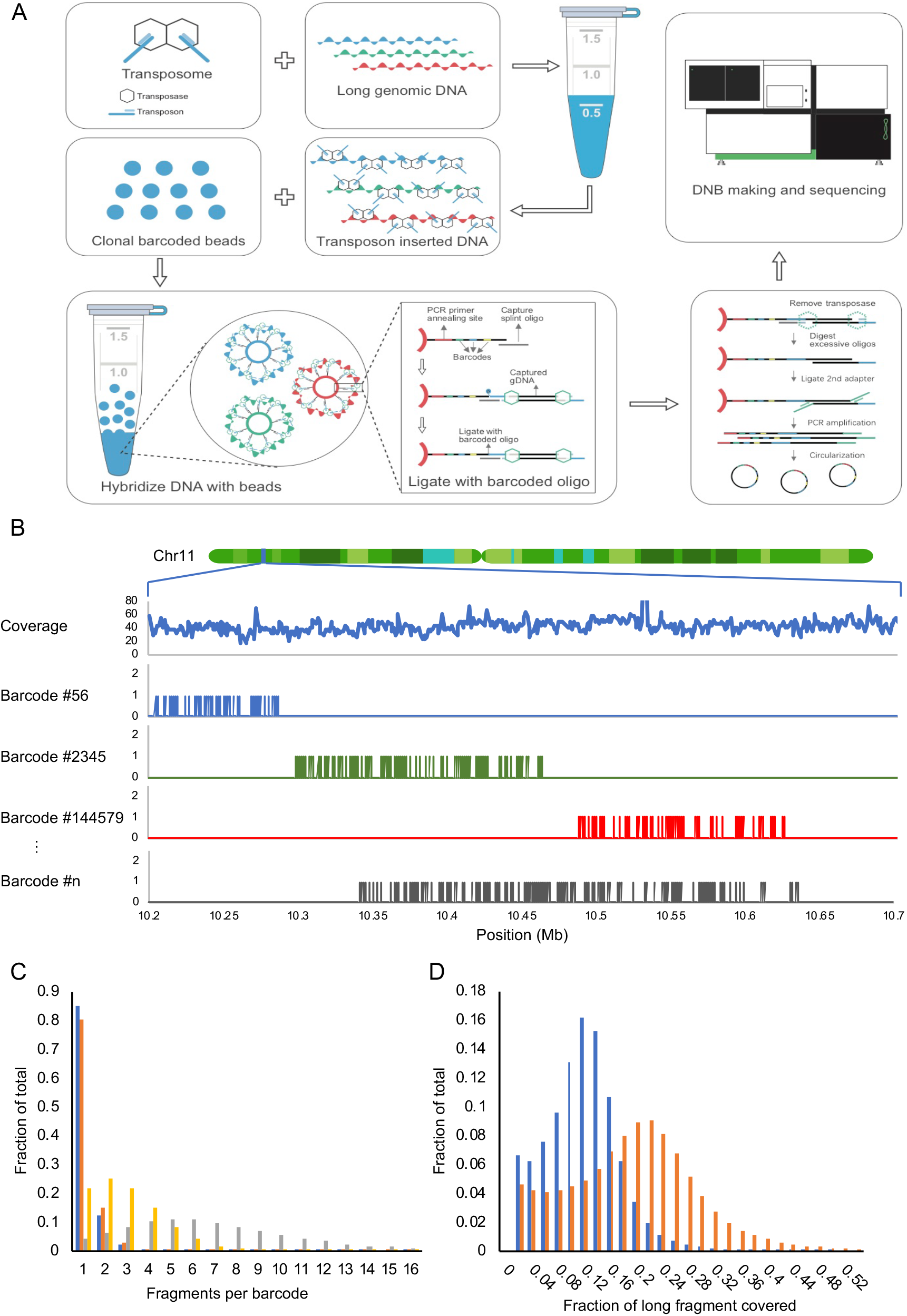
Overview of stLFR. (A) The first step of stLFR involves inserting a hybridization sequence approximately every 200-1000 base pairs on long genomic DNA molecules. This is achieved using transposons. The transposon integrated DNA is then mixed with beads that each contain ~400,000 copies of an adapter sequence that contains a unique barcode shared by all adapters on the bead, a common PCR primer site, and a common capture sequence that is complementary to the sequence on the integrated transposons. After the genomic DNA is captured to the beads, the transposons are ligated to the barcode adapters. There are a few additional library processing steps and then the co-barcoded sub-fragments are sequenced on a BGISEQ-500 or equivalent sequencer. (B) Mapping read data by barcode results in clustering of reads within 10 to 350 kb regions of the genome. Total coverage and barcode coverage from 4 barcodes are shown for the 1 ng stLFR-1 library across a small region on Chr11. Most barcodes are associated with only one read cluster in the genome. (C) The number of original long DNA fragments per barcode are plotted for the 1 ng libraries stLFR-1 (blue) and stLFR-2 (orange) and the 10 ng stLFR libraries stLFR-3 (yellow) and stLFR-4 (grey). Over 80% of the fragments from the 1 ng stLFR libraries are co-barcoded by a single unique barcode. (D) The fraction of nonoverlapping sequence reads (blue) and captured sub-fragments (orange) covering each original long DNA fragment are plotted for the 1 ng stLFR-1 library.

In one approach to stLFR, two different transposons are used in the initial insertion step, allowing PCR to be performed after exonuclease treatment. However, this approach results in approximately 50% less coverage per long DNA molecule as it requires that two different transposons were inserted next to each other to generate a proper PCR product. To achieve the highest coverage per genomic DNA fragment we use a single transposon in the initial insertion step and add an additional adapter through ligation. This noncanonical ligation, termed 3’ branch ligation, involves the covalent joining of the 5’ phosphate from the blunt-end adapter to the recessed 3’ hydroxyl of the genomic DNA (Figure 1a). A detailed explanation of this process has previously been described (Wang et al. 2018). Using this method, it is theoretically possible to amplify and sequence all sub-fragments of a captured genomic molecule. In addition, this ligation step enables a sample barcode to be placed adjacent to the genomic sequence for sample multiplexing. This is useful as it does not require an additional sequencing primer to read this barcode. After the ligation step, PCR is performed and the library is ready to enter any standard next generation sequencing (NGS) workflow. In the case of BGISEQ-500, the library is circularized as previously described (Drmanac et al. 2010). From single stranded circles DNA nanoballs are made and loaded onto patterned nanoarrays (Drmanac et al. 2010). These nanoarrays are then subjected to combinatorial probe-anchor synthesis (cPAS) based sequencing on the BGISEQ-500 (Fehlmann et al. 2016; Huang et al. 2017; Mak et al. 2017). After sequencing, barcode sequences are extracted using a custom program (Materials and Methods). Mapping the read data by unique barcode shows that most reads with the same barcode are clustered in a region of the genome corresponding to the length of DNA used during library preparation (Figure 1b). A protocol of this process and the process used to make clonally barcoded beads has been described by Cheng *et al.* (Cheng et al. 2018).

### stLFR read coverage and variant calling

To demonstrate stLFR phasing and variant calling we generated four libraries using 1 ng (stLFR-1 and stLFR-2) and 10 ngs (stLFR-3 and stLFR-4) of DNA isolated from cell line GM12878. The number of beads added ranged from 10 million (stLFR-4), 30 million (stLFR-3), and 50 million (stLFR-1 and stLFR-2). Finally, the 3’ branch ligation method was used for libraries stLFR-1-3 and the two-transposon method was used for stLFR-4. Both stLFR-1 and stLFR-2 were sequenced deeply to 336 Gb and 660 Gb of total base coverage, respectively. We also analyzed these at downsampled coverages. stLFR-3 and stLFR-4 were sequenced to more modest levels of 117 Gb and 126 Gb, respectively. Co-barcoded reads were mapped to build 37 of the human reference genome using BWA-MEM (Li and Durbin 2009). The non-duplicate coverage ranged from 34-58X and the number of long DNA molecules per barcode ranged from 1.2-6.8 (Table 2 and Figure 1c). As expected, the stLFR libraries made from 50 million beads and 1 ng of genomic DNA had the highest single unique barcode co-barcoding rates of up to 85% (Figure 1c). These libraries also observed the highest average non-overlapping read coverage per long DNA molecule of 10.7-12.1% and the highest average non-overlapping base coverage of captured sub-fragments per long DNA molecule of 17.9-18.4% (Figure 1d). This coverage is ~10X higher than previously demonstrated using 3 ng of DNA and transposons attached to beads (Zhang et al. 2017). This suggests our solution-based transposition process is 3-fold more efficient at sub-fragment capture (40.7-47.4 sub-fragments per genomic fragment in 1 ng of genomic DNA versus 5 sub-fragments captured in 3 ng at similar read coverage as reported by Zhang et al. (Zhang et al. 2017).

**Table 2.**
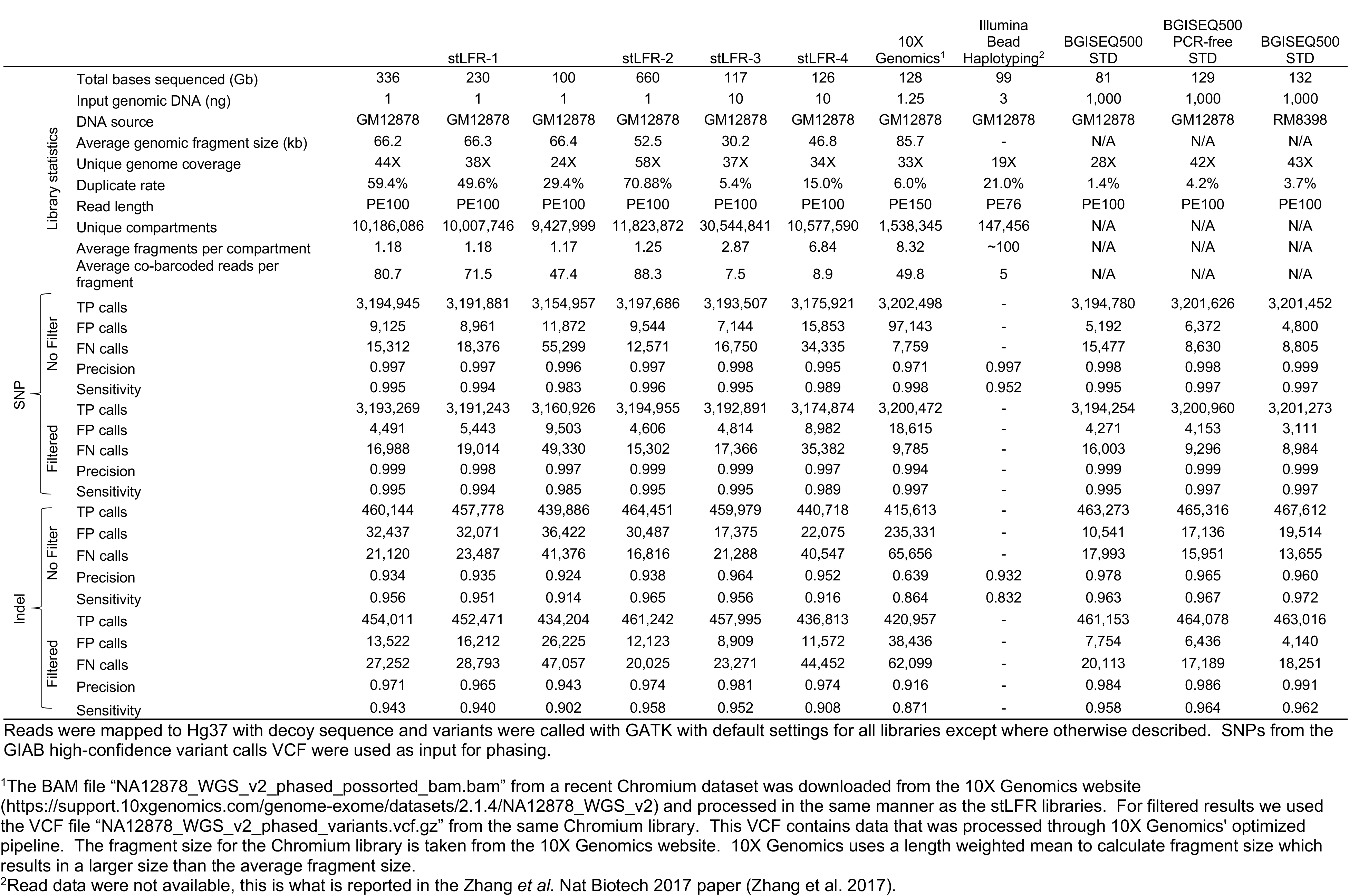
Variant calling statistics.

For each library variants were called using Sentieon’s DNAseq (Freed et al. 2017) using default settings. Comparing SNP and indel calls to Genome in a Bottle (GIAB) (Zook et al. 2014) allowed for the determination of true positive (TP), false positive (FP), and false negative (FN) rates (Table 2). In addition, we performed variant calling using the same settings on standard non-stLFR libraries made from GIAB reference material (NIST RM 8398) DNA and the same genomic DNA used to make stLFR libraries. We also compared precision and sensitivity rates to a Chromium library made by 10X Genomics (Zheng et al. 2016) and to those reported in the bead haplotyping library study by *Zhang et al.* (Zhang et al. 2017). Although direct comparisons can be difficult due to differences in coverage, for most metrics of variant calling stLFR libraries performed similar or better than the published results from Zhang et al. or Chromium libraries, especially when nonoptimized mapping and variant calling processes were used (Table 2, “No Filter”). To further improve the variant calling performance in stLFR libraries we used a machine learning algorithm trained against additional stLFR libraries made from GIAB samples GM12878, GM24385, GM24149, GM24143, and GM24631 (Supplemental Table 1 and Methods). This lead to the discovery of a few selection criteria that lowered the FP rate by ~40%. Importantly, this was achieved while increasing the FN rate by less than 10% in most stLFR libraries. Taking into account these variants and the reduced number of FP variants after filtering results in a similar FP rate and a 2-3 fold higher FN rate than the filtered STD library for SNP calling (Table 2).

One potential issue with using GIAB data to measure the FP rate is that we were unable to use the GIAB reference material (NIST RM 8398) due to the rather small fragment size of the isolated DNA. For this reason, we used the GM12878 cell line and isolated DNA using a dialysis-based method capable of yielding very high molecular weight DNA (see methods). However, it is possible that our isolate of the GM12878 cell line could have a number of unique somatic mutations compared to the GIAB reference material and thus cause the number of FPs to be inflated in our stLFR libraries. To examine this further single nucleotide FP variants were compared across all of the NA12878 libraries (Supplemental Fig. 1a). 1,740 FP variants were shared between stLFR libraries 1-4 and both standard libraries made from GM12878 cell line DNA, but not shared with the standard library made from GIAB reference material. We also compared cell line DNA FPs with the Chromium library and found that 1,268 of these shared FPs were also present in the Chromium library (Supplemental Fig. 2b). Importantly, the Chromium library was sequenced using Illumina technology. Examination of the distribution of these shared FP variants across the genome versus randomly selected true positive variants (Supplemental Fig. 2) showed similar patterns with the vast majority of shared FP variants having distributions similar to randomly selected variants. 60 shared FP variants were found within 100 bases of each other and could be the result of incorrect mapping due to short insert sizes. This suggests that the stLFR process introduces very few FP errors, this is likely due to the low (~1,000X) amount of amplification used to make these libraries.

### stLFR phasing performance

We developed a custom software program called LongHap (Methods) to make full use of the unique characteristics of stLFR data. Filtered variants called by each respective library were used for phasing within that library. In general, phasing performance was very high with over 99% of all heterozygous SNPs in most samples placed into contigs with N50s ranging from 1.2-34.0 Mb depending on the library type and the amount of sequence data (Supplemental Table 2). Comparison to GIAB data showed that short and long switch error rates were low (Supplemental Table 2) and comparable to previous studies (Mao et al. 2016; Zheng et al. 2016; Zhang et al. 2017). The stLFR-1 library with 336 Gb of total read coverage (44X unique genome coverage) achieved the highest phasing performance with a contig N50 of 34.0 Mb (Figure 2). Indeed, the entire length of most chromosomes were covered by only a few contigs (Figure 2). Importantly, even with only 100 Gb of data (24X unique genome coverage) the phased contig N50 was still 14.4 Mb (Figure 2). N50 length appeared to be mostly affected by length and coverage of long genomic fragments. This can be seen in the decreased N50 of stLFR-2 as the DNA used for this sample was slightly older and more fragmented than the material used for stLFR-1 (average fragment length of 52.5 kb versus 62.2 kb) and the ~10-fold shorter N50s of stLFR-3 and stLFR-4 made from 10 ngs of DNA (Supplemental Table 2).

**Figure 2.**
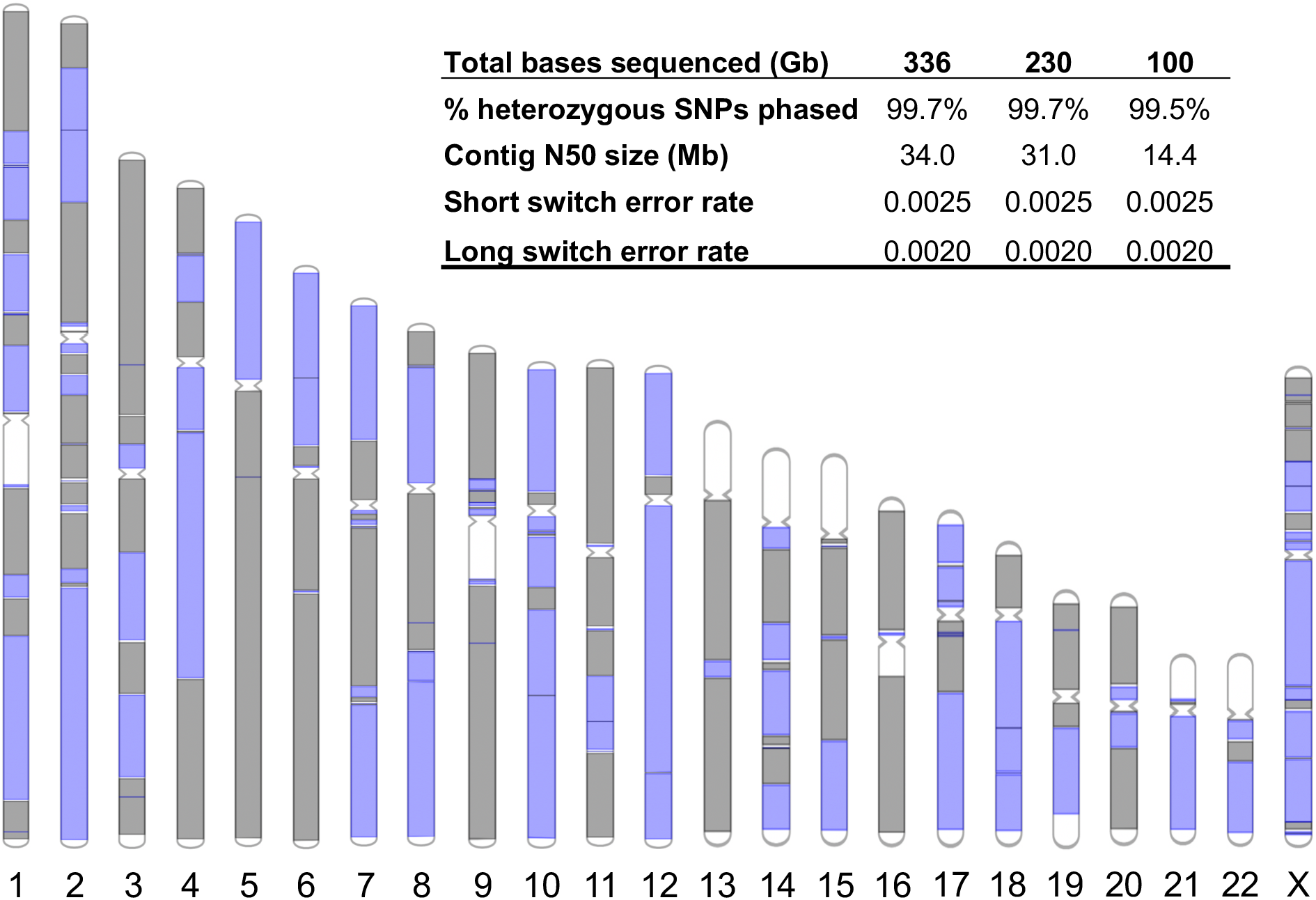
stLFR-1 phasing performance. The 221 phased blocks from the stLFR-1 library are depicted on chromosomes as alternating colors of grey and purple. Unphased regions are depicted in white. The inset table shows the performance of phasing with different sequence read coverage levels.

**Figure 3.**
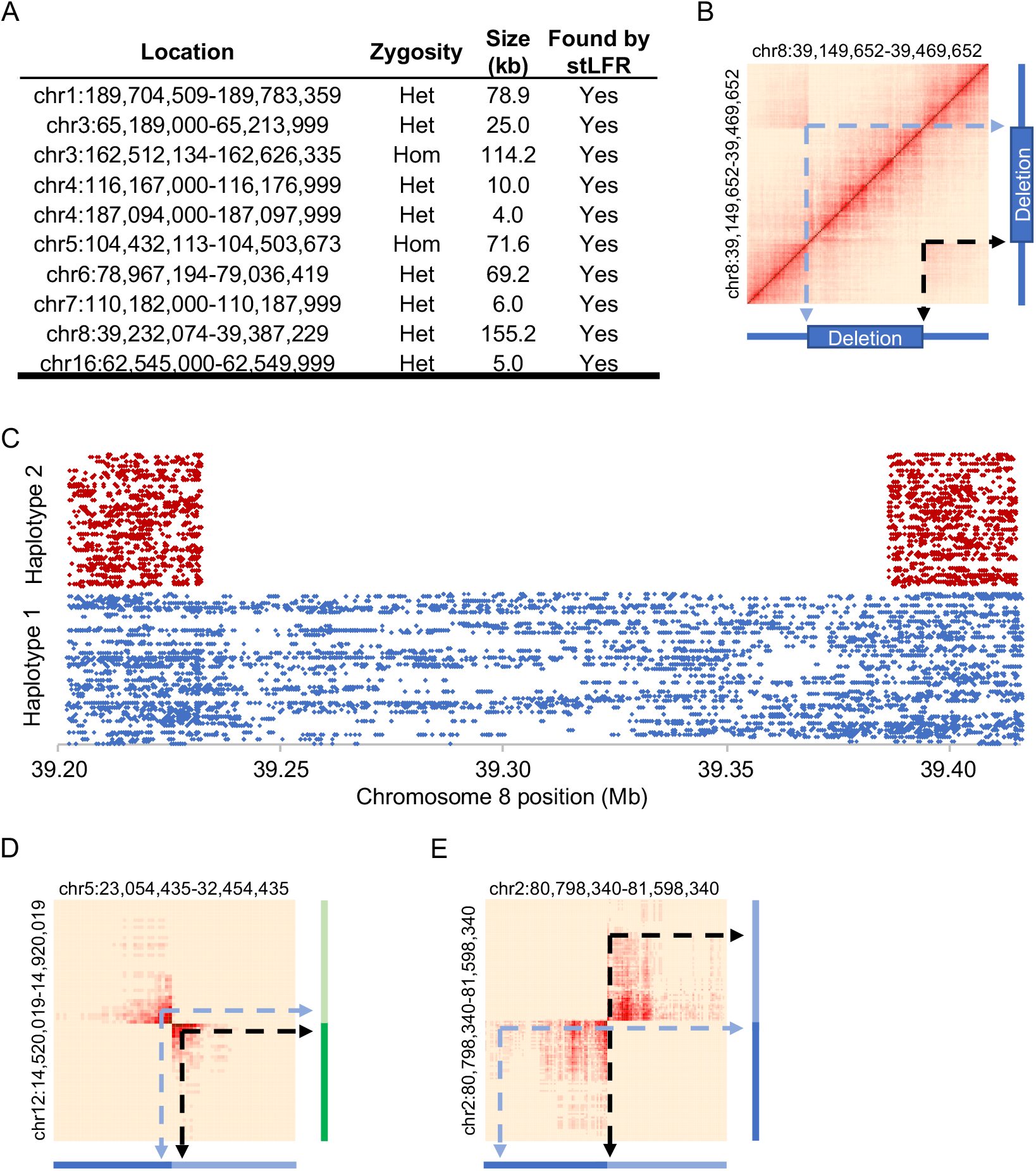
SV detection. (A) Previously reported deletions in NA12878 were also found using stLFR data. Heat maps of barcode sharing for each deletion can be found in Supplemental Fig. 3. (B) A heat map of barcode sharing within windows of 2kb for a region with a ~150 kb heterozygous deletion on chromosome 8 was plotted using a Jaccard Index as previously described (Zhang et al. 2017). Regions of high overlap are depicted in dark red. Those with no overlap in beige. Arrows demonstrate how regions that are spatially distant from each other on chromosome 8 have increased overlap marking the locations of the deletion. (C) Co-barcoded reads are separated by haplotype and plotted by unique barcode on the y axis and chromosome 8 position on the x axis. The heterozygous deletion is found in a single haplotype. (D) Heat maps were also plotted for overlapping barcodes between chromosomes 5 and 12 for a patient cell line with a known translocation (Dong et al. 2016) and (E) GM20759, a cell line with a known transversion in chromosome 2 (Dong et al. 2017).

In an effort to make a fair comparison of stLFR phasing performance to a Chromium library (the Zhang *et al.* bead haplotyping method did not have read data available making direct comparisons impossible) we attempted to phase only the high confidence variant call set from GIAB. This removed any influence of the variant calling performance within a library from the phasing performance of that library. We also used HapCut2 (Edge et al. 2017), a freely available software package designed for phasing co-barcoded and Hi-C sequencing data. Overall, the phasing performance of stLFR was similar to the Chromium library. The number of SNPs phased and the long and short switch error rates were essentially identical (Supplemental Table 2). Similar to the results from LongHap, stLFR-1 generated the longest contig N50 of 15.1 Mb, which was similar to the N50 achieved by the Chromium library. In all cases it should be noted that conclusions of performance differences are difficult due to differences in input DNA length, total read coverage, and sequencing platforms used.

### Structural variation detection

Previous studies have shown that long fragment information can improve the detection of structural variations (SVs) and described large deletions (4-155 kb) in NA12878 (Zheng et al. 2016; Zhang et al. 2017). To demonstrate the power of stLFR to detect SVs we examined barcode overlap data, as previously described (Zhang et al. 2017), for stLFR-1 and stLFR-4 libraries in these regions. In every case the deletion was observed in the stLFR-1 data, even at lower coverage (Figure 2a and Supplemental Fig. 6). Closer examination of the co-barcoded sequence reads covering a ~150 kb deletion in chromosome 8 demonstrated that the deletion was heterozygous and found in a single haplotype (Figure 2b-c). The 10 ng stLFR-4 library also detected most of the deletions, but the three smallest were difficult to identify due to the lower coverage per fragment (and thus less barcode overlap) of this library.

To evaluate stLFR performance for detecting other types of SVs we made libraries from a cell line from a patient with a known translocation between chromosomes 5 and 12 (Dong et al. 2016) and GM20759, a cell line with a known inversion on chromosome 2 (Dong et al. 2017). stLFR libraries were able to identify the inversion and the translocation in the respective cell lines (Figure 2d-e). Downsampling the amount of reads per library showed that a strong signal of the translocations was detected even with as little as 5 Gb of read data (~1.7X total coverage, Supplemental Fig. 4a-h). Finally, examination of both SVs in the stLFR-1 library resulted in no obvious pattern (Supplemental Fig. 4i-l), suggesting the false positive rate for detection of these types of SVs is low.

### *de novo* assembly with stLFR

For 1 ng input libraries up to 85% of fragments were co-barcoded by a single unique barcode. This means the majority of barcodes should be associated with reads derived from very small regions of the genome (<300 kb). This type of information is akin to single molecule long sequencing data and should enable diploid *de novo* genome assembly. To test if stLFR can be used for *de novo* assembly we used stLFR-1 and stLFR-2 libraries and the software package Supernova 2.1.1 (10X Genomics, Pleasanton, CA). This software was not designed for stLFR and as a result does not allow for data with over 4.7 million barcodes to be used. Due to this limitation, we had to reduce the total number of barcodes for each stLFR library by combining over 10 million barcodes into a total of 4.7 million barcodes for this analysis. This is not ideal as it reduces the amount of information, but remarkably stLFR data performed very well with this software package. Contig and scaffold N50s of ~100 kb and ~30 Mb, respectively, were achieved for both libraries (Table 3). Plotting the assembled contigs against chromosome sequences from Genome Reference Consortium Human Build 38 (GRCh38) showed high concordance (Figure 4). Analysis of the resulting assemblies with the Quality Assessment Tool for Genome Assemblies (QUAST) (Gurevich et al. 2013) and comparison to other assemblies of NA12878 using Chromium (Zheng et al. 2016) or nanopore (Jain et al. 2018) technologies suggested that the stLFR derived assemblies were very complete and harbored few misassembled regions (Supplemental Table 3).

**Figure 4.**
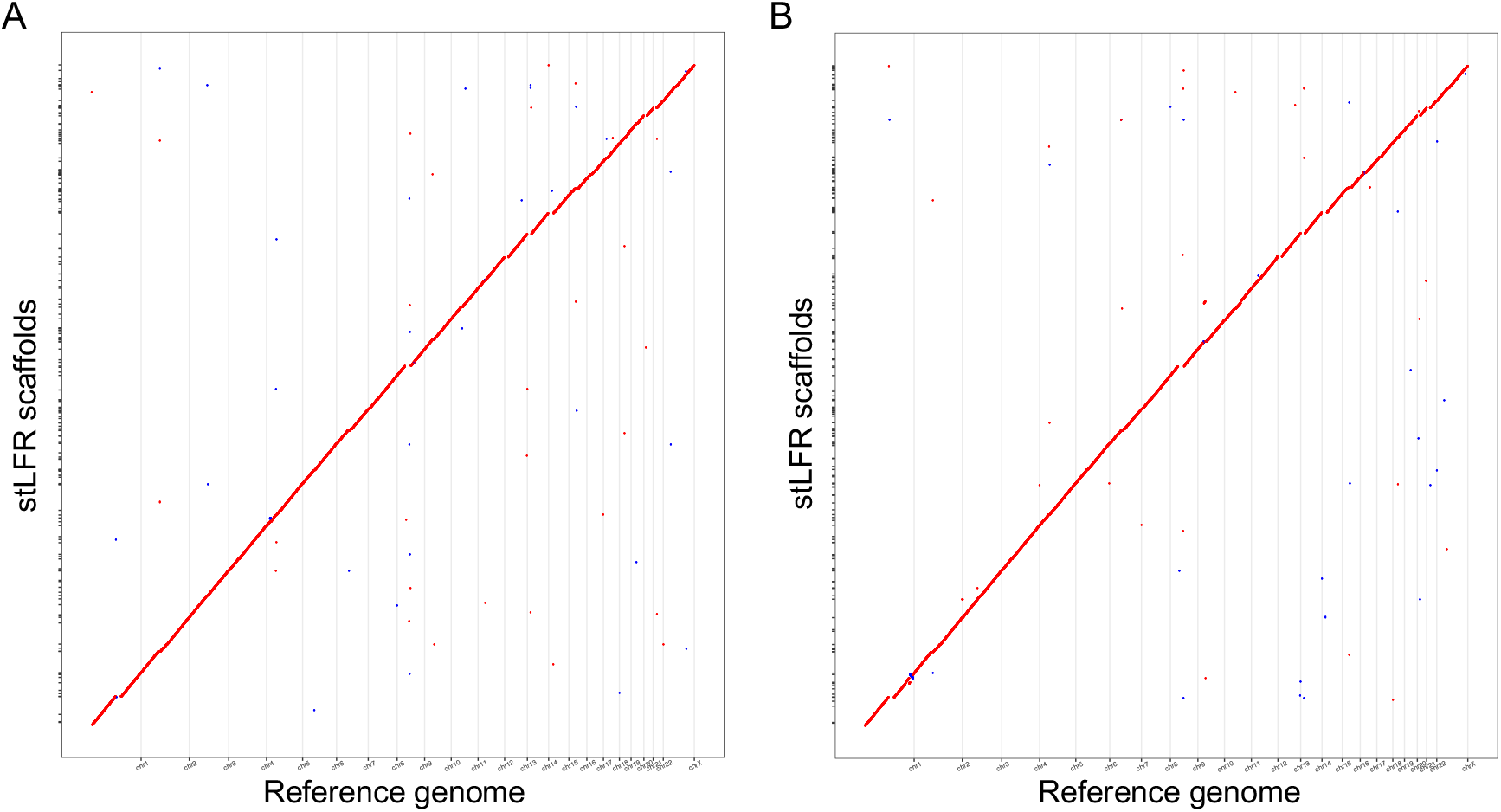
Dot plots of *de novo* assembled NA12878. The scaffolds from the *de novo* assemblies of stLFR-1 (A) and stLFR-2 (B) were compared against chromosomes from GRCh38 using dot plots.

**Table 3.** NA12878 *de novo* assembly statistics.

|  | stLFR-1 | stLFR-2 |
| --- | --- | --- |
| Contig N50 (kb) | 99.78 | 99.73 |
| Scaffold N50 (Mb) | 29.02 | 29.65 |
| Phase Block N50 (Mb) | 2.2 | 1.43 |
| Assembly Size (Gb) | 2.73 | 2.72 |

## Discussion

Here we describe an efficient whole genome sequencing library preparation technology, stLFR, that enables the co-barcoding of sub-fragments of long genomic DNA molecules with a single unique clonal barcode in a single tube process. Using microbeads as miniaturized virtual compartments allows a practically unlimited number of clonal barcodes to be used per sample at a negligible cost. Our optimized hybridization-based capture of transposon inserted DNA on beads, combined with 3’-branch ligation and exonuclease degradation of the excess capture adapters, successfully barcodes up to ~20% of sub-fragments in DNA molecules as long as 300 kb in length. Importantly, this is achieved without DNA amplification of initial long DNA fragments and the representation bias that comes with it. In this way, stLFR solves the cost and limited co-barcoding capacity of emulsion-based methods.

The quality of variant calls using stLFR is very high and possibly, with further optimization, will approach that of standard WGS methods, but with the added benefit that co-barcoding enables advanced informatics applications. Using stLFR we demonstrate high quality, near complete phasing of the genome into long contigs with extremely low error rates, detection of SVs, and *de novo* assembly of a human genome. All of this is achieved from a single library that does not require special equipment nor additional library preparation costs.

As a result of efficient barcoding, we successfully used as little as 1 ng of human DNA (600 X genome coverage counting top and bottom strands of each DNA molecule) to make stLFR libraries and achieved high quality WGS with most sub-fragments uniquely co-barcoded. Less DNA can be used, but stLFR does not use DNA amplification during co-barcoding and thus does not create overlapping sub-fragments from each individual long DNA molecule. For this reason overall genomic coverage suffers as the amount of DNA is lowered. In addition, a sampling problem is created as stLFR currently retains 10-20% of each original long DNA molecule followed by PCR amplification. This results in a relatively high sequencing read duplication rate and additional sequencing cost. One potential solution is to remove the PCR step. This would eliminate sampling, but also it could substantially reduce the false positive and false negative error rates. In addition, improvements such as optimizing the distance of insertion between transposons and increasing the length of sequencing reads to paired-end 200 bases are relatively easy to enable and would increase the coverage and overall quality. For some applications, such as structural variation detection, using less DNA and less coverage may be desirable. As we demonstrate in this paper, as little as 5 Gb of sequence coverage can faithfully detect inter and intrachromosomal translocations and in these cases the duplication rate is negligible. Indeed, stLFR may represent a simple and cost-effective replacement for long mate pair libraries in a clinical setting.

In addition, we believe this type of data will enable routine near perfect diploid phased *de novo* assembly from a single stLFR library without the need for long contiguous reads such as those generated by SMRT or nanopore technologies. One interesting feature of transposon insertion is that it creates a 9 base sequence overlap between adjacent sub-fragments. Frequently, these neighboring sub-fragments are captured and sequenced enabling reads to be synthetically doubled in length (e.g., for 200 base reads, two neighboring captured sub-fragments would create two 200 base reads with a 9 base overlap, or 391 bases).

In this paper libraries were made with 50 million beads, however using more is possible. This will enable many types of cost-effective analyses where 100s of millions of barcodes would be useful. We envision this type of cheap massive barcoding can be useful for RNA analyses such as full-length mRNA sequencing from 1,000s of cells by combination with single cell technologies or deep population sequencing of 16S RNA in microbial samples. Phased chromatin mapping by the Assay for Transposase-Accessible Chromatin (ATAC-seq) (Buenrostro et al. 2013) or methylation studies are all also possible with stLFR. Finally, in an effort to share what we believe to be a very important technology, we have made a detailed protocol freely available for academic use (Cheng et al. 2018).

## Materials and Methods

### High Molecular Weight DNA isolation

Long genomic DNA was isolated from cell lines following a modified version of the RecoverEase™ DNA isolation kit (Agilent Technologies, La Jolla, CA) protocol (Agent Technologies 2015). Briefly, approximately 1 million cells were pelleted and lysed with 500 ul of lysis buffer. After a 10 minute incubation at 4 °C 20 μL of RNase-IT ribonuclease cocktail in 4 mL of digestion buffer was added directly to the lysed cells and incubated on a 50 °C heat block. After 5 minutes 4.5 mL of proteinase K solution (~1.1 mg/mL proteinase K, 0.56% SDS, and 0.89X TE) was added and the mix was incubated at 50 °C for an additional 2 hours. The genomic DNA was then transferred to dialysis tubing with a 1,000 kD molecular weight cutoff (Spectrum Laboratories, Inc., Rancho Dominguez, CA) and dialyzed overnight at room temperature in 0.5X TE buffer.

### Barcoded bead construction

Barcoded beads are constructed through a split and pool ligation-based strategy using three sets of double-stranded barcode DNA molecules. A common adapter sequence was attached to Dynabeads^™^ M-280 Streptavidin (ThermoFisher, Waltham, MA) magnetic beads with a 5’ dual-biotin linker. Three sets 1,536 of barcode oligos containing regions of overlapping sequence were constructed by Integrated DNA Technologies (Coralville, IA). Ligations were performed in 384 well plates in a 15 μL reaction containing 50 mM Tris-HCl (pH 7.5), 10 mM MgCl_2_, 1 mM ATP, 2.5% PEG-8000, 571 units T4 ligase, 580 pmol of barcode oligo, and 65 million M-280 beads. Ligation reactions were incubated for 1 hour at room temperature on a rotator. Between ligations beads were pooled into a single vessel through centrifugation, collected to the side of the vessel using magnet, and washed once with high salt wash buffer (50 mM Tris-HCl (pH 7.5), 500 mM NaCl, 0.1 mM EDTA, and 0.05% Tween 20) and twice with low salt wash buffer (50 mM Tris-HCl (pH 7.5), 150 mM NaCl, and 0.05% Tween 20). Beads were resuspended in 1X ligation buffer and distributed across 384 wells plates and the ligation steps were repeated.

### stLFR using two transposons

2 pmol of Tn5 coupled transposons were inserted into 40 ng of genomic DNA in a 60 μL reaction of 10 mM TAPS-NaOH (pH 8.5), 5 mM MgCl_2_, and 10% DMF at 55 °C for 10 minutes. 1.5 μL of transposon inserted DNA was transferred to 248.5 μL of hybridization buffer consisting of 50 mM Tris-HCl (pH 7.5), 100 mM MgCl_2_, and 0.05% TWEEN^®^ 20. 10-50 million barcoded beads were resuspended in the same hybridization buffer. The diluted DNA was added to the barcoded beads and the mix was heated to 60 °C for 10 minutes with occasional light mixing. The DNA-bead mix was transferred to a tube revolver in a laboratory oven and incubated at 45 °C for 50 minutes. 500 uL of ligation mix containing 50 mM Tris-HCl (pH 7.8), 10 mM DTT, 1 mM ATP, 2.5% PEG-8000, and 4,000 units of T4 ligase was added directly to the DNA-bead mix. The ligation reaction was incubated at room temperature on a revolver for 1 hour. 110 μL of 1% SDS were added and the mix was incubated at room temperature for 10 minutes to remove the Tn5 enzyme. Beads were collected to the side of the tube via a magnet and washed once with low salt wash buffer and once with NEB2 buffer (New England Biolabs, Ipswich, MA). Excess barcode oligos were removed using 10 units of UDG (New England Biolabs, Ipswich, MA), 30 units of APE1 (New England Biolabs, Ipswich, MA), and 40 units of Exonuclease 1 (New England Biolabs, Ipswich, MA) in 100 uL of 1X NEB2 buffer. This reaction was incubated at 37 °C for 30 minutes. Beads were collected to the side of the tube and washed once with low salt wash buffer and once with 1X PCR buffer (1X PfuCx buffer (Agilent Technologies, La Jolla, CA), 5% DMSO, 1 M Betaine, 6 mM MgSO_4_, and 600 μM dNTPs). The PCR mix containing 1X PCR buffer, 400 pmol of each primer, and 6 μL of PfuCx enzyme (Agilent Technologies, La Jolla, CA) was heated to 95 °C for 3 minutes then cooled to room temperature. This mix was used to resuspend beads and the combined mixture was incubated at 72 °C for 10 minutes followed by 12 cycles of 95 °C for 10 seconds, 58 °C for 30 seconds, and 72 °C for 2 minutes.

### stLFR with 3’ branched adapter ligation

This method starts with the same hybridization insertion conditions but using only one transposon as opposed to two transposons. After capture and barcode ligation steps, as described above, beads were collected to the side of the tube and washed with low salt wash buffer. An adapter digestion mix of 90 units of Exonuclease I (New England Biolabs, Ipswich, MA) and 100 units of Exonuclease III (New England Biolabs, Ipswich, MA) in 100 μL of 1X TA Buffer (Teknova, Hollister, CA) is added to the beads and incubated at 37 °C for 10 minutes. The reaction is stopped and the Tn5 enzyme is removed by adding 11 μL of 1% SDS. Beads were collected to the side of the tube and washed once with low salt wash buffer and once with 1X NEB2 buffer (New England Biolabs, Ipswich, MA). Excess capture oligo was removed by adding 10 units of UDG (New England Biolabs, Ipswich, MA) and 30 units of APE1 (New England Biolabs, Ipswich, MA) in 100 uL of 1X NEB2 buffer (New England Biolabs, Ipswich, MA) and incubating at 37 °C for 30 minutes. Beads were collected to the side of the tube and washed once with high salt wash buffer and once with low salt wash buffer. 300 pmol of second adapter was ligated to the bead bound sub-fragments with 4,000 units of T4 ligase in 100 uL of ligase buffer containing 50 mM Tris-HCl (pH 7.8), 10 mM MgCl_2_, 0.5 mM DTT, 1 mM ATP, and 10% PEG-8000 on a revolver for 2 hours at room temperature. Beads were collected to the side of the tube and washed once in high salt wash buffer and once in 1X PCR buffer. The PCR mix and conditions were the same as the two-transposon process described above.

### Sequence mapping and variant calling

Raw read data were first demultiplexed by the associated barcode sequence using the barcode split tool (available at GitHub https://github.com/stLFR/stLFR_read_demux). Barcode assigned and clipped reads were mapped to the hs37d5 reference genome with BWA-MEM (Li and Durbin 2009). The resulting BAM file was then sorted by chromosomal coordinates with SAMtools (Li et al. 2009) and duplicates were marked with Picard MarkDuplicate function (http://broadinstitute.github.io/picard). Short variant (SNPs and indels) calling was performed using Sentieon’s DNAseq (Freed et al. 2017) optimization of the GATK’s HaplotypeCaller (McKenna et al. 2010). To further improve the FP rate in stLFR libraries, we developed a binary classification model for variant filtering based on XGboost (Chen and Guestrin 2016). TPs and FPs from samples GM12878, GM24385, GM24149, GM24143, and GM24631 (Supplemental Table 1) were generated using VCFeval (Cleary et al. 2015) by comparing to Genome in a Bottle (GIAB) high confidence variant lists (Zook et al. 2014) for each sample and labeled for model training. Using custom software (https://github.com/stLFR/extremevariantfilter) mapping quality (MQ), MQ rank sum, strand odds ratio, fisher strand bias, read position rank sum, quality by depth, reference allele depth, alternate allele depth, percentage of reads supporting the reference allele, the ratio of alternate depth to reference depth, and an encoding of genotype (homozygous or heterozygous) were extracted from the labeled VCFs and used as features for model training. Models were trained individually for SNPs and InDels and generalizability of the models was tested by training on four of the five samples and testing on the fifth.

### LongHap

We developed a novel phasing algorithm, LongHap (https://github.com/stLFR/stLFR_LongHap), specifically designed for the uniqueness of stLFR’s data. A seed-extension strategy was employed in the phasing process. It initially starts from one pair of seeds, composed of the most upstream heterozygous variant in the chromosome. The seeds are extended by linking the other downstream candidate variants until no more variants can be added to the extending seeds (Supplemental Fig. 5). In this extending process, the candidate variants at different loci will not be equally treated (*i.e.*, the upstream variant has higher priority compared with the downstream ones across the chromosome). Each two heterozygous loci have two possible combinations along the two different alleles. Taking variant T_2_/G_2_ and G_3_/C_3_ for example (Supplemental Fig. 5), one combination pattern is T_2_-G_3_ and G_2_-C_3_, while another one is T_2_-C_3_ and G_2_-G_3_. The score of each combination is calculated by the number of long DNA fragments spanning the two loci, which is equivalent to the number of unique barcodes with reads mapping to these two loci. As shown in Supplemental Fig. 5, the final score of the former combination is 3, which is three times more than the latter. The variant T_2_/G_2_ is added to the extending seeds and the process repeats. Notably, if any barcode supports both of the alleles at one specific locus, it will be ignored when calculating the linkage score. This helps to decrease the switch error rate. When a conflict in linking downstream candidate variants occurs, as the variant A_4_/C_4_ in Supplemental Fig. 5 shows, a simple decision will be made by comparing the linked loci number to allow further extending candidate variants. In this case, there are two linked loci in the left scenario while there is only one in the right scenario. LongHap will choose the left combination pattern as the final phasing result.

### Variant phasing with Hapcut2

SNPs were phased with Hapcut2 (https://github.com/vibansal/HapCUT2) (Edge et al. 2017) using its 10X Genomics data pipeline. The BAM file was first converted into a format that carries barcode information in a similar format as a 10X Genomics barcoded BAM. Specifically, a ‘BX’ field was added to each line reflecting the barcode information of that read. GIAB variants or variants called by GATK for each library were used as the input for phasing, and the phasing result was summarized and compared against the GIAB phased vcf file (Zook et al. 2014) using the calculate_haplotype_statistics.py tool of Hapcut2.

### SV detection

Structural variants were detected by calculating shared barcodes between regions of the genome as previously described (Zhang et al. 2017). Duplicate reads were first removed. The mapped co-barcoded reads were scanned using a sliding window (the default value is 2 kb) along the genome, every window recorded how many barcodes have been found within this 2 kb window, and a Jaccard index was calculated for the shared barcodes ratio between the window pairs. Structural variant events were identified by the Jaccard index sharing metric between window pairs.

For every window pair (X, Y) across the genome, the Jaccard index is calculated as follows:

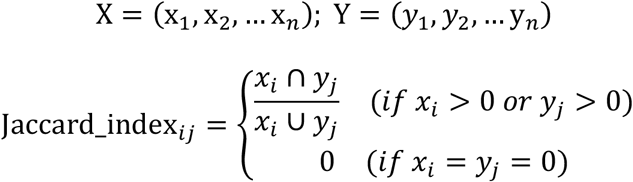

### *De novo* assembly

For each library, barcodes with a minimum of 10 reads were selected, and these barcodes were degenerated into the list of ~4.7 million barcodes from the Longranger2.2.2 software package (10X Genomics, Pleasanton, CA) barcode whitelist (located in unpacked software at /longranger-2.2.2/longranger-cs/2.2.2/tenkit/lib/python/tenkit/barcodes/4M-with-alts-february-2016.txt). stLFR fastq files were converted into a format that resembles Chromium fastq files and were then used as input for Supernova 2.0.1 (10X Genomics, Pleasanton, CA). Specifically, for supernova runs, options maxreads = 2100000000 and nopreflight were used. A pseudohap assembly output was generated with the mkoutput function of Supernova and scaffolds with a minimum length of 10 kb were compared against the human reference GRCh38 with QUAST 5.0.0 (Gurevich et al. 2013). In addition, pseudohap2 assembly outputs were also generated from supernova, and each haplotype was aligned to GRCh38 with function nucmer of mummer 4.0.0 (Delcher et al. 1999; Kurtz et al. 2004), with options -c 1000 -l 100. The alignment delta file was filtered with the delta-filter function of mummer to filter for one-to-one alignments. Scaffolds of at least 500 kb and alignments of at least 50 kb from the resulting alignment list were plotted into dotplots.

## Acknowledgments

We would like to acknowledge the ongoing contributions and support of all Complete Genomics and BGI-Shenzhen employees, in particular the many highly skilled individuals that work in the libraries, reagents, and sequencing groups that make it possible to generate high quality whole genome data. We would also like to thank Z. Dong, Z. Yang, and W. Xie for providing cell lines for the translocation analysis. This work was supported in part by the Shenzhen Peacock Plan (NO.KQTD20150330171505310) and the National Key Research and Development Program of China (NO.2017YFC0906501). B.A.P. is a recipient of and this work was partially supported by the Research Fund for International Young Scientists, National Natural Science Foundation of China (31550110216).

## Authors contributions

R.D. and B.A.P. conceived the study. O.W., R.C., X.C., M.K.W., H.K.L., D.C., L.W., F.F., Y.Z., S.D., D.N., A.A., X.X., R.D., and B.A.P. developed the molecular biology process of stLFR. R.Y.Z., S.D., S.G., N.B., and A.C. performed the sequencing. Q.M., J.T., Y.S., Y.Z., E.A., Y.X., C.V., S.N., W.T., J.W., X.L., X.Q., H.W., Y.D., and Z.L. developed algorithms for and performed analyses on stLFR data. O.W., C.X., J.S.L., W.Z., H.Y., J.W., K.K., X.X., R.D., and B.A.P. coordinated the study. O.W., R.D., and B.A.P. wrote the manuscript. All authors reviewed and edited the manuscript.

## Completing interests

Employees of BGI and Complete Genomics have stock holdings in BGI.

## Data access

All sequencing data reported in this paper have been deposited in the China National GeneBank (CNGB) Nucleotide Sequence Archive (CNSA, db.cngb.org/cnsa/) under accession number CNP0000066.

## Supplementary Materials

Figures S1-S5

Tables S1-S3

